# Chimpanzees and bonobos reinstate an interrupted triadic game

**DOI:** 10.1101/2023.10.05.560857

**Authors:** Raphaela Heesen, Adrian Bangerter, Klaus Zuberbühler, Katia Iglesias, Federico Rossano, Jean-Pascal Guéry, Emilie Genty

## Abstract

When humans engage in joint action, they seem to so with an underlying sense of joint commitment, a feeling of mutual obligation towards their partner and a shared goal. Whether our closest living relatives, bonobos and chimpanzees, experience and understand joint commitment in the same way is subject to debate. Crucial evidence concerns how participants respond to interruptions of joint actions, particularly if they protest or attempt to reengage their reluctant or distracted partners. During dyadic interactions, bonobos and chimpanzees appear to have some sense of joint commitment, according to recent studies. Yet, data are inconsistent for triadic games with objects. We addressed this issue by engaging *N*=23 apes (5 adult chimpanzees, 5 infant bonobos, 13 adult bonobos) in a “tug-of-war” game with a human experimenter who abruptly stopped playing. Adult apes readily attempted to reengage the experimenter (>60% of subjects on first trial), with no group differences in the way of reengagement. Infant bonobos rarely reengaged and never did so on their first trial. Importantly, when infants reengaged passive partners, they mostly deployed (tactile) signals, yet rarely game-related behaviours (GRBs) as commonly observed in adults. These findings might explain negative results of earlier research. Bonobos and chimpanzees may thus have motivational foundations for joint commitment, although this capacity might develop over lifetime. We discuss this finding in relation to evolutionary and developmental theories on joint commitment.

## 1. Introduction

Many social animals engage in collaborative activities where two or more participants work towards a goal that would be unreachable by individuals. For instance, chimpanzees, whales, hyenas, and fish engage in group hunting [1–4], ants cooperatively transport objects [5,6], and meerkats mob predators [7]. Humans though engage in *joint actions* with a presumably different underlying psychology which enables them to ‘share’ their intentions [8,9]. Shared intentionality, it has been argued, may form the basis of many cultural achievements, social institutions, and language [10]. Specifically, sharing intentions encompasses the formation of *joint commitment* – a construct that describes a feeling of mutual obligation towards the partner to bring a joint action to completion, and which supposedly underpins all human collaborative activities [9,11–13].

Empirically, joint commitment appears to manifest in specific behavioural patterns [14,15], although there is some debate on whether behavioural markers can truly represent mental constructs as such [16]. The presumed behavioural correlates include ostensive signalling upon entering, maintaining, and exiting joint actions [14,15,17–21] and, when faced with interruptions, attempts to reinstate the joint action in a coordinated fashion [22–24]. Humans in some cultural settings perceive sudden, not mutually ratified, interruptions as socially inappropriate [19,25], which typically leads to negative emotional reactions and corrective actions towards the partner [26–29]. It is noteworthy however that the current definition and empirical findings derive from observations and experiments conducted in Western, Educated, Industrial, Rich and Democratic (WEIRD) populations; and it thus remains to be demonstrated whether joint commitment represents a universal human construct.

Interruptions of joint activities nonetheless provide a means to comparatively investigate joint commitment. One classic paradigm consists of an experimenter abruptly disengaging from triadic games involving objects, such as to bounce a woodened block on a trampoline by holding it on opposite sites [26,27,30]. Moreover, Gräfenhain and colleagues [26] tested two- to three-year-old children’s reactions to an interruption initiated by an experimenter in either a no-commitment condition (i.e., child and experimenter each play on their own) or a joint commitment condition (i.e., they play the game together). Three-year-olds, more so than two-year-olds, attempted to reengage partners significantly more often in the joint than the no commitment condition, suggesting a developmental trajectory of joint commitment. Young children thus already appear to understand that dissolving a commitment requires prior mutual agreement [28]. Thus, understanding of joint commitment seems to emerge gradually, starting around the age of three and becoming more profound at the age of five years, alongside a more general awareness of shared intentions and social norms [31].

Investigations of whether non-human animals, notably our closest relatives, the great apes, can experience something akin to joint commitment have led to inconclusive results. While some researchers claim that joint commitment is human-unique [10,30], contrasting evidence on how great apes coordinate joint actions, both naturally and in experimental situations, have reopened discussions [20,24,32–37]. Despite this new evidence, joint commitment continues to be claimed a uniquely human capacity, primarily because of negative evidence in chimpanzees [30] as well as still too-limited experimental evidence [38]. The argument largely rests on one influential comparative study on human children and three young chimpanzees, aged between 33 and 51 months [30]. Here, the children, but none of the chimpanzees, attempted to reengage the reluctant human experimenter, which was taken as evidence of joint commitment in humans yet not in apes. Reengagement in human children was interpreted as an attempt to repair with others the breakdown of a joint “we” [31]. However, subsequent studies testing older apes between 3-7 years and in less complex social interactions came to different conclusions, both concerning interactions with human experimenters [34,36,39] as well as joint activities with conspecifics [24,32,37]. Moreover, considering newer reports on reengagement in bonobos [24,36], species differences in terms of reengagement behaviour could equally explain negative findings in chimpanzees [30]. Potential differences between bonobos and chimpanzees might be explained by differential social styles. Bonobos have a reputation of being less despotic and more tolerant than chimpanzees [40–42], though this pattern varies between groups and settings [e.g., 43].

In this study, we used a triadic game paradigm to look for evidence of reengagement attempts in chimpanzees and bonobos, and different age classes in bonobos. Specifically, we implemented a “tug of war” game between a human experimenter and the apes. The game started by pulling a garden hose back and forth. After a few iterations, the experimenter suddenly stopped by letting go of the hose. If apes had a potential sense of joint commitment, we predicted that they would attempt to reengage the human experimenter, for instance by using game-related behaviours “GRBs” (e.g., handing back the object) or other communicative signals. To address potential age effects, we compared reengagement behaviours of infant and adult bonobos, expecting infants to be less likely to reengage than adults given fewer experiences in coordinating joint activities, notably with objects. We further manipulated the experimenter’s attentional state, by either looking towards (“still-faced condition”) or away from the subject (“back-turned condition”), as an additional source of variation in reengagement behaviour. We expected subjects to be more likely to attempt resuming the game when experimenters gazed at the subject, compared to when being turned away, as gaze might be interpreted as a signal of availability or readiness for interaction. This assumption is based on research in apes and human children, which showed that gaze represents a potential coordination device, with eye-contact being understood as a signal to engage [20,44,45].

## 2. Materials & Methods

### 2.1 Study Groups

#### 2.1.1 Infant Bonobos

Data on five infant bonobos (*Pan paniscus*, mean age = 3.4 y; SD = 1.0 y; range = 2.5-4.5 y; 2 females; 3 males, see S1 Table) were collected from October 2018 until November 2018, at the Lola Ya Bonobo sanctuary in the Democratic Republic of Congo. The infants were orphans, mostly victims of the bushmeat trade and confiscated by authorities. They had been cared for by humans from the moment of their arrival at the sanctuary. Each infant was cared for by a human surrogate mother for a few years before being introduced into an existing social group. During the day, they could range freely in an outdoor enclosure of approximately 500 m^2^, comprising a forested patch that offered climbing opportunities. In addition, the enclosure was equipped with climbing structures, ropes, a pool, and a trampoline. The infants were free to interact with other orphans or the human surrogate mothers, who were always present in the enclosure. The infants were bottle-fed with a mixture of cow milk and cereals with water twice a day, and additionally received fruits, sugar cane, peanuts, and vegetables. Each individual received 5 l of water per day. On rainy days, an indoor enclosure was available (approximately 150 m^2^), provided with climbing structures and ropes and the testing isolation cage (5 m^2^). At night they slept in the indoor enclosures (each cage sized about 2 m^2^, 2 infants per cage), which were furnished with hammocks.

#### 2.1.2 Adult Bonobos

Our experiment included thirteen subadult and adult bonobos, hereafter referred to as “adult bonobos” (mean age = 13.5 y; SD = 8.2 y; age range: 6.0 - 30.0 y; 9 females; 4 males; see S1 Table), also housed at the Lola Ya Bonobo sanctuary (i.e., data also collected from October 2018 until November 2018). The bonobos lived in three different social groups: Group 1: 20 individuals, including 11 females; Group 2: 19 individuals, including 8 females; Group 3: 15 individuals, including 8 females. The groups inhabited three 8-15 ha enclosures, consisting of streams, swamps, lakes, primary rainforest, and grassy open areas. The enclosures were separated by fence. They were provided with food by caregivers in four feedings per day (6-9 types of vegetables per day at 9 am, 2-4 types of fruits per day at 11 am and again at 4 pm, and protein balls at 2 pm), but could also range freely within their enclosures to forage for herbaceous vegetation and wild fruits. Water was available from streams, lakes, and keepers (provided through bottles during the day). During the night, the groups were held in 75 m^2^ dormitories furnished with hammocks.

#### 2.1.3 Adult Chimpanzees

At the time of study, the chimpanzee group consisted of seven chimpanzees (*Pan troglodytes verus)*, of which two individuals did not participate in the study (two females, aged 10 and 11 years). Thus, we tested five adult chimpanzees (mean age = 19.2 y; SD = 5.76 y; range: 9.0 - 23.0 y; 1 female; 4 males; see S1 Table), housed at the zoological park of La Vallée des Singes, France. The chimpanzees were tested between May and October 2018. The chimpanzee facility included an outside enclosure with a large forest area and climbing structures in grassy areas (7,500 m^2^), and an inside enclosure with enrichment and various climbing structures (220 m^2^). In stable weather conditions, the group was kept outside. Food was distributed five to six times a day and includes daily rations of primate pellets, fruits, vegetables, and rice. The chimpanzees were also occasionally fed with nuts, meat once a week, and eggs twice a week, and can forage for natural vegetation in their outdoor enclosure (wild berries, herbaceous vegetation). Fresh water was always available from a source at the building and a stream surrounding the island.

### 2.2 Experimental Design and Procedure

We tested subjects using a tug-of-war game by which a human experimenter and an ape pulled on opposite ends of a plastic garden hose through the mesh or bars of their indoor or outdoor compartments (adult bonobos, Fig 1b), through the bars of their indoor lodges (adult chimpanzees, Fig 1b) and from within the inside of the testing cage (infant bonobos; Fig 1a). For the game to work, i.e., to obtain repeated sequences of back-and-forth pulling movements of the hose, both partners had to alternate pulling the hose, and the apes (who are much stronger than humans) had to adjust their pulling force, otherwise the hose would be pulled inside the cage and the game would stop. The chimpanzees had been in contact with garden hoses prior to the experiment, either as part of the cleaning equipment of keepers or as a toy inside the cage. For the bonobos, it was uncertain whether they had any experience with garden hoses; thus, we implemented a habituation period by distributing 2-3 garden hoses identical to those used in the experiment in all enclosures one week before the start of the experiment. Participation was voluntary, meaning that during testing no adult subject was isolated from the rest of their social group and could come or leave as they pleased. Infant bonobos on the other hand were isolated from the other orphans during testing to avoid regular disturbances by other playful infants (see section 2.2.1 for details); during testing, the orphans’ surrogate human mother was present in the cage for emotional support.

**Fig 1.**
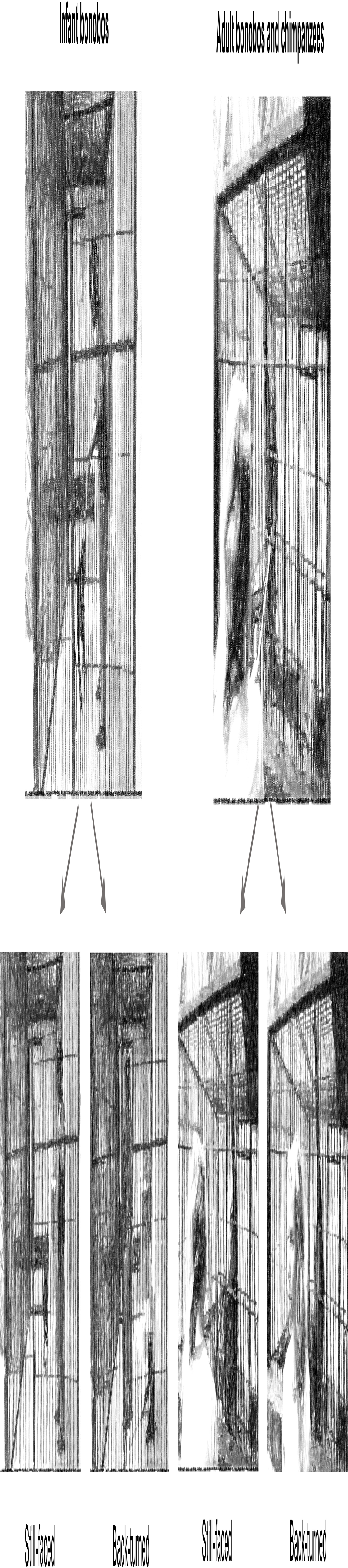
Experimental set-up for infant bonobos (a) and for adult bonobos and chimpanzees. **(b).** When testing infant bonobos, the experimenter remains inside of the cage and engages in a tug-of-war game with the subject in the cage, using a garden hose. When testing adults, the experimenter stands outside the cage and engages in a tug-of-war game with the subject through the cage mesh, also using a garden hose. In the still-faced condition, the experimenter interrupts the game and faces the subject while remaining still; in the back-turned condition, the experimenter interrupts the game and turns their back to the subject while remaining still. Raphaela Heesen provided consent to be shown in the images linked to (b).

All sessions were filmed from outside the cage using a Panasonic HC-V770 Camcorder on a tripod, equipped with an external Sennheiser microphone (MKE 400). Trials began either by an ape subject handing the hose to the experimenter or by the experimenter handing the hose to the subject, accompanied by verbal encouragement. As soon as both held the hose, the goal was to establish a minimum of three rapid back-and-forth pulling movements (“tug-of-war”). This movement was then suddenly interrupted by the experimenter who dropped the hose (for interruption period durations, see sections below). During an interruption, the experimenter stood still, either facing the subject (still-faced condition) or turning their back to the subject (back-turned condition), see Fig 1. Two additional conditions were included in the original study plan but could not be implemented in a sufficiently consistent fashion across sites to be included in the study (see S1 Text).

Trials and conditions were applied opportunistically given that the subjects’ individual motivation to participate in the game varied across testing days. Consequently, each experimental session could include one or more trials. Each of the 4 initial conditions (still-faced and back-turned, as well as the two failed conditions, see S1 Text) were presented in a randomized order, and could be administered once or several times depending on the subject’s motivation. As much as possible, we tried to counterbalance the order of conditions across testing days. We tried to test all subjects at least once in each condition, but as this depended largely on their motivation, not every individual was tested in each condition.

#### 2.2.1 Infant Bonobos

Infant bonobos were tested in the isolation cage of their indoor enclosure (Fig 1a). Experimenters were the infants’ human surrogate mothers, to whom they are emotionally attached (i.e., besides of a few trials conducted by author RH). The subjects were carried to the testing compartment by their surrogate mothers who entered the cage with them. The hose was placed in the cage before they arrived. Once they entered the testing cage, the door was locked during the session. Each testing session was stopped after 15 min even if no trial had been completed; if the infants were motivated to play, the session could be longer. For each session and subject, experimenters were selected depending on their availabilities during the day, thus the order in which infants were tested was based on the mother’s most convenient schedule. On a testing day, subjects were called into the testing area and brought in by their surrogate mother; if they were not motivated, they were called in again later, or testing was postponed to another day. Following a previous experiment with bonobos with a similar paradigm [46], the interruption periods lasted 30 s. We analysed all behaviours and communicative signals occurring during this period. Surrogate mothers had reported no prior experience in conducting the tug-of-war game with infant bonobos. In total, we were able to conduct 29 trials with infant bonobos (still-faced: *N* = 17; *median* [*IQR*] = 3 [2; 4] trials per individual; back-turned: *N* = 12; *median* [*IQR*] = 2 [2; 3] trials per individual).

#### 2.2.2 Adult Bonobos

The subjects were tested in their indoor and outdoor facility, wherever the experimenter and the subjects could play the game through the cage mesh. For group 3 and 1, this was in primarily in their isolation cage or indoors, e.g., Fig 1b. For group 2, this was outside either in the isolation cage or through the mesh doors of the outdoor enclosure. Four different persons acted as experimenters: two women (authors RH and EG who interacted rarely with the bonobos), and two men (zookeepers working full-time with the bonobos). Like for infants, the interruption periods also lasted 30 s, during which we investigated all behaviours and communicative signals. The session lasted as long as the bonobos were motivated to participate. For adult bonobos (and chimpanzees), no session cap of 15 min was applied; the subjects were not isolated from the rest of their group like infant bonobos but could come and go as they pleased (i.e., opposite to the orphans, the testing did not restrict the adult subjects’ abilities to engage in other daily social activities). All testing was voluntary, and we recruited participants based on their motivation to play with the experimenter. Keepers and researchers had reported no prior experience in conducting the tug-of-war game with the adult bonobos. The only exception is one mother-reared individual (Moyi, “MO”, S1 Table) who had engaged in a tug-of-war like game with a cotton rag with author EG in July 2013, an event that initially sparked the idea to use such a game for this experiment. We conducted 52 trials with adult bonobos (still-faced: *N* = 30; *median* [*IQR*] = 1 [1; 4] trials per individual; back-turned: *N* = 22; *median* [*IQR*] = 1 [1; 2.5] trials per individual).

#### 2.2.3 Adult Chimpanzees

The game was played inside the holding building, and through the cage mesh, at a location wherever subjects spontaneously engaged in the game, but always in the indoor enclosures either in the mornings or evenings. Five persons acted as experimenters. These included two women (author RH who interacted rarely with the chimpanzees and a zookeeper who worked approx. once a month with the chimpanzees) and three men (one zookeeper working approx. five times a month with the chimpanzees, and another who worked full-time with chimpanzees). The keepers’ participation in a testing session was determined based on their availabilities on a given testing day. To allow comparison with chimpanzees in the study by Warneken et al. [30], the interruption periods were aimed at lasting 15 s. However, in some cases the keepers failed to react immediately to the researcher’s cue to interrupt the game, resulting in interruption periods that slightly varied in length (lasting on average 22.1 s, *SD* = 7.76 s). To allow for a more consistent evaluation of all behaviors and communicative signals between trials, we therefore only analyzed behaviors or communicative signals occurring during the first 15 s of the interruption period. As for the other groups, the session lasted as long as the chimpanzees were motivated to participate in the game. Keepers and researchers had no prior experience in conducting the tug-of-war game with the chimpanzees. We were able to conduct 58 trials with adult chimpanzees (still-faced: *N* = 27; *median* [*IQR*] = 5 [3; 6] trials per individual; back-turned: *N* = 31; *median* [*IQR*] = 3 [3; 4] trials per individual).

### 2.3 Video Coding

We coded GRBs and signals (i.e., gestures, vocalizations, facial expressions) deployed during interruption periods using the ELAN package, version 5.2 [47]. During interruption periods, we annotated whether subjects attempted to reengage their partner or not. A reengagement attempt was annotated (yes = 1/no = 0) if one or several GRBs or communicative signals were produced during an interruption (see Table 1 and ethogram in S3 Table).

**Table 1.**
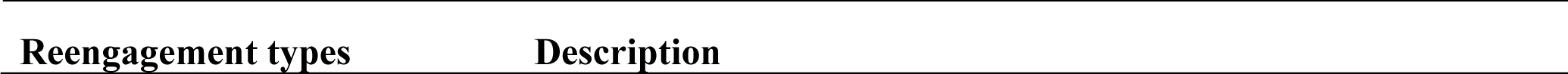
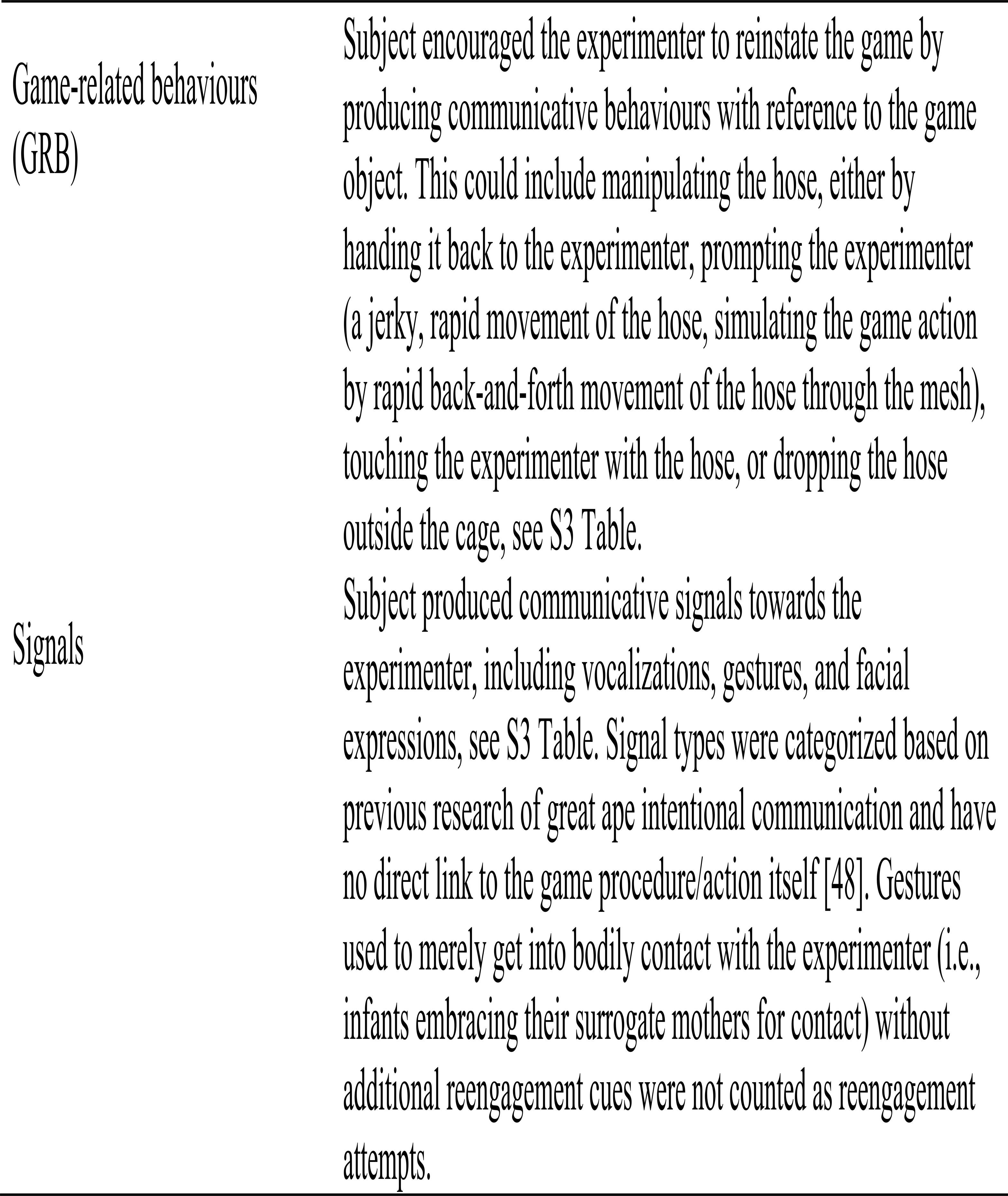
Communicative behaviours indicating reengagement during interruption periods.

#### 2.3.1 Coding Reliability

We carried out an inter-observer reliability test between two great ape communication researchers (authors RH and EG). The test compared the coding on the presence or absence of GRBs and signals (i.e., which indicated the presence or absence of reengagement), based on the ethogram in S3 Table, occurring during the interruption periods in both the still-faced and back-turned conditions. Double coding was done for a total of 22 trials, representing 15 % of trials for each group out of their total number of trials as reported in the main text. The result for signals was Cohen’s κ = 0.81 (agreement 98.9 %) and for GRB Cohen’s κ = 0.84 (agreement 95.5 %), indicating almost perfect agreement [49].

### 2.4 Statistical Analysis

We first present results on the number of subject that reengaged in each group, both considering only the first and subsequent trials (i.e., overall), as well as the number of subjects that used GRBs/signals on first and subsequent trials (yielding Table 3).

Additionally, as subsequent analysis, we considered reengagement, GRBs and signal percentages based on repeated measures and variation in the number of trials across subjects. We computed percentages of reengagement attempts (number of “yes”) by dividing each participant’s number of reengagement attempts by its number of trials, multiplied by 100. To describe the results, we presented median percentage of these reengagement attempts, as well as the interquartile range (IQR), i.e., first and third quartile. Likewise, for signals and GRBs, we computed the percentage of reengagement attempts in which subjects used signals and GRBs by dividing the number of reengagements attempts *with* signal or GRBs by the number of subjects’ total of *reengaged* trials; in our results, we presented the median of these percentages along with the IQRs.

We further tested the effect of condition (still-faced and back-turned condition *within* group), age (comparison infants and adult bonobos) and group (comparison adult chimpanzees and adult bonobos) on the percentages by which individuals produced a) reengagement attempts, b) GRBs, and c) signals as per reengaged trials. We used non-parametric Wilcoxon signed-rank tests for related comparisons within group and Mann-Whitney U tests (i.e., Wilcoxon’s rank-sum test) for independent between-group comparisons. Following [50], we report results for the Wilcoxon signed-rank tests via *p*-values, and results for the Mann-Whitney U tests via the test statistic provided in R (*W*) as well as respective *p-value*s. To indicate the magnitude of existing effects, we additionally computed effect sizes (“*r*”) for between-group comparisons that were significant at *alpha =* 0.05 following the R function provided in [50]. Effects exceeding the threshold of 0.3 indicate medium effects (accounting for 9% of the total variance) and effects exceeding 0.5 indicate large effects (accounting for 25% of the variance), see [50]. Subjects who only had participated in one condition were excluded from the analysis of the Wilcoxon signed-rank tests. Where sample sizes were too small (e.g., when comparing GRB and signal use between groups *and* within condition), we reported the combined results for both conditions (i.e., sections 3.4 and 3.5).

To test whether the differences in experimental design between chimpanzees and bonobos could have affected our results, we decided to conduct additional posthoc analyses (section 3.6). Since chimpanzees’ and bonobos’ interruption periods differed in duration (i.e., 15 sec and 30 sec, respectively), we conducted a posthoc group comparison between adult bonobos and chimpanzees, but in which we only considered reengagement attempts within the first 15 s.

Moreover, our experiment was affected by subjects’ level of motivation, insofar as each subject participated voluntarily without being forced or separated from their social group. This yielded an unwanted variation in the number of trials across subjects, which may have affected our results on reengagement rates. To take account of this, we conducted a Spearman’s rank correlation test to examine the relationship between number of trials and reengagement percentages.

Lastly, since some of our adult bonobo subjects are orphans and were raised by human surrogate mothers rather than by their natural mothers (*N* = 10, see S1 Table), early experiences through human interactions could have fostered reengagement behaviour. Therefore, the subjects of this group cannot be directly compared without additional verification of rearing impact. Thus, we provide additional results on reengagement separately for orphans and non-orphan adult bonobos (see section 3.2.1, adult bonobos).

### 2.5 Ethics statement

We received ethical agreement for this study from the Commission d’Ethique de la Recherche of the University of Neuchâtel (agreement number: 01-FS-2017), the internal ethical committee of La Vallée des Singes, and from the Ministère de la Recherche Scientifique et Technologie de la République Démocratique du Congo (research permit number: MIN.RST/SG/180/020/2018). Raphaela Heesen provided consent to be shown in the image of Fig 1.

## 3. Results

Descriptive results on latencies of reengagement responses across groups are presented in Table 2.

**Table 2.**
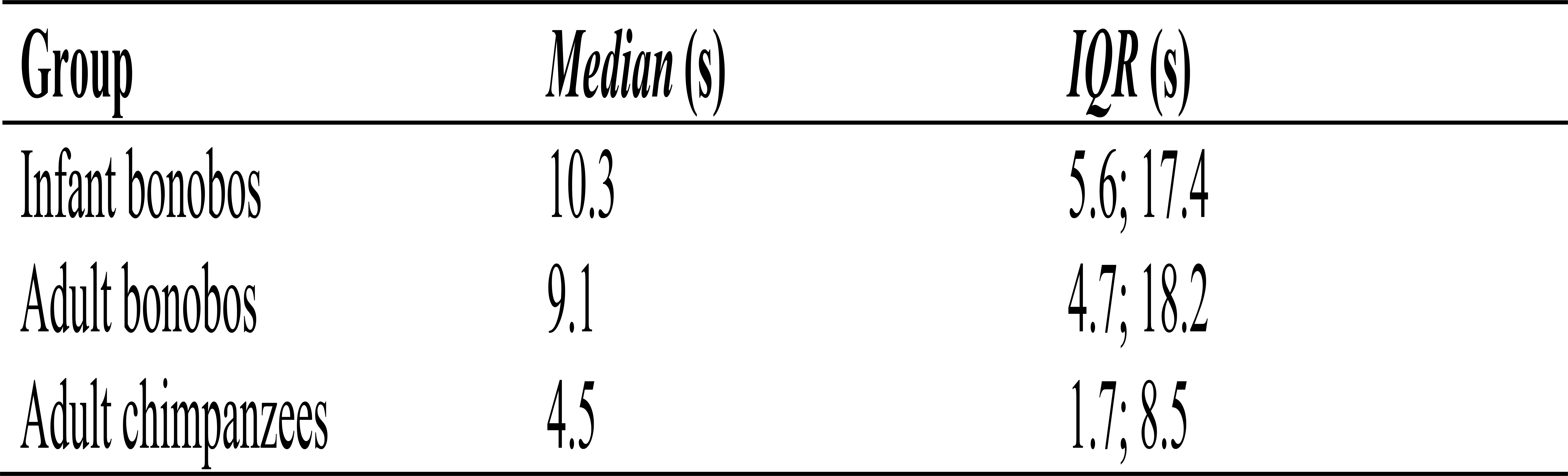
Response latencies (s) of reengagement attempts across groups.

### 3.1 Infant bonobos

#### 3.1.1 Do infant bonobos reengage passive partners?

On first trial, none of the infant bonobos reengaged the experimenters (0%, Table 3). When considering both first and subsequent trials, four out of five infant bonobos reengaged the experimenter in at least in one trial (80%, Table 3).

**Table 3.**
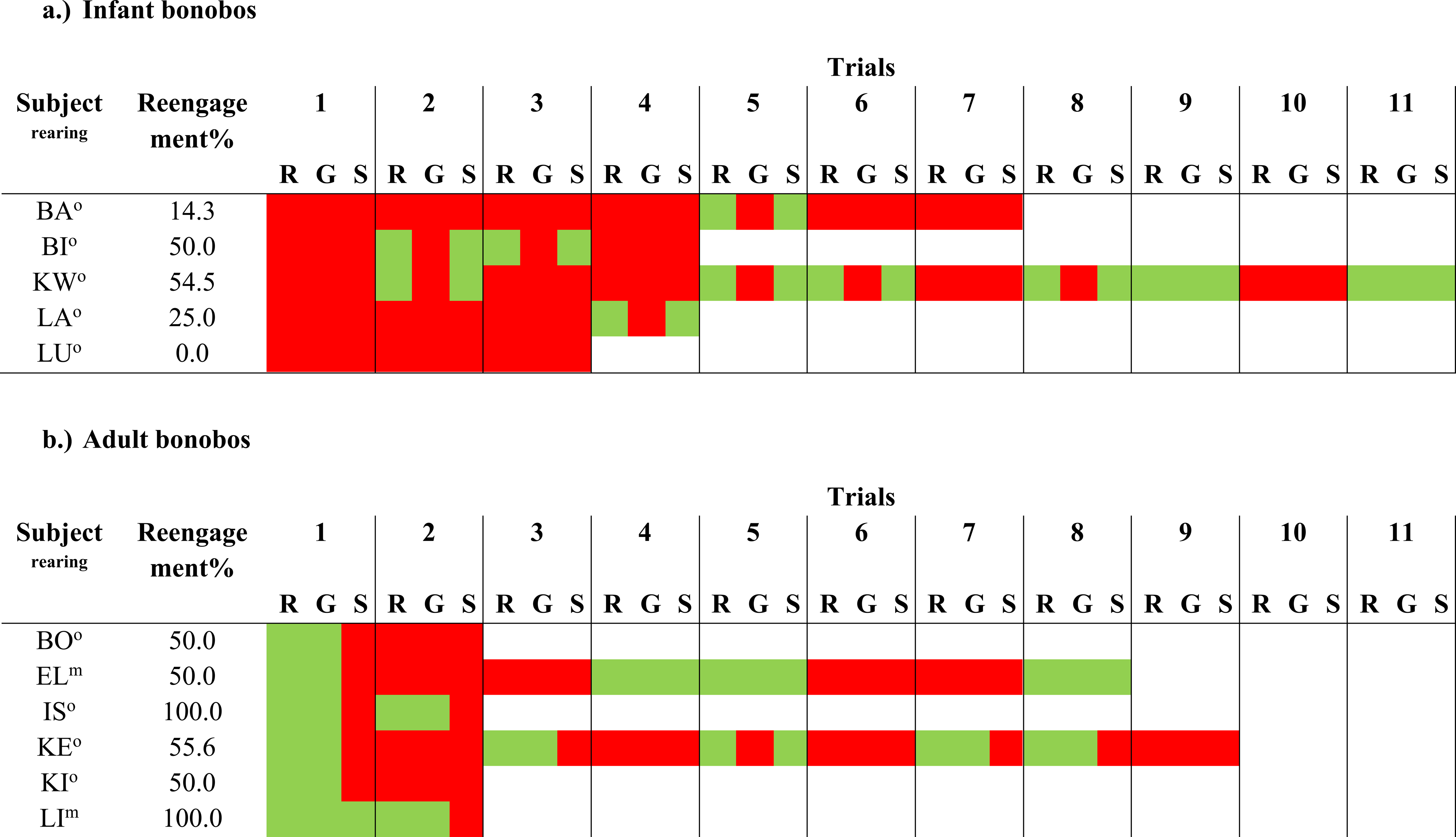
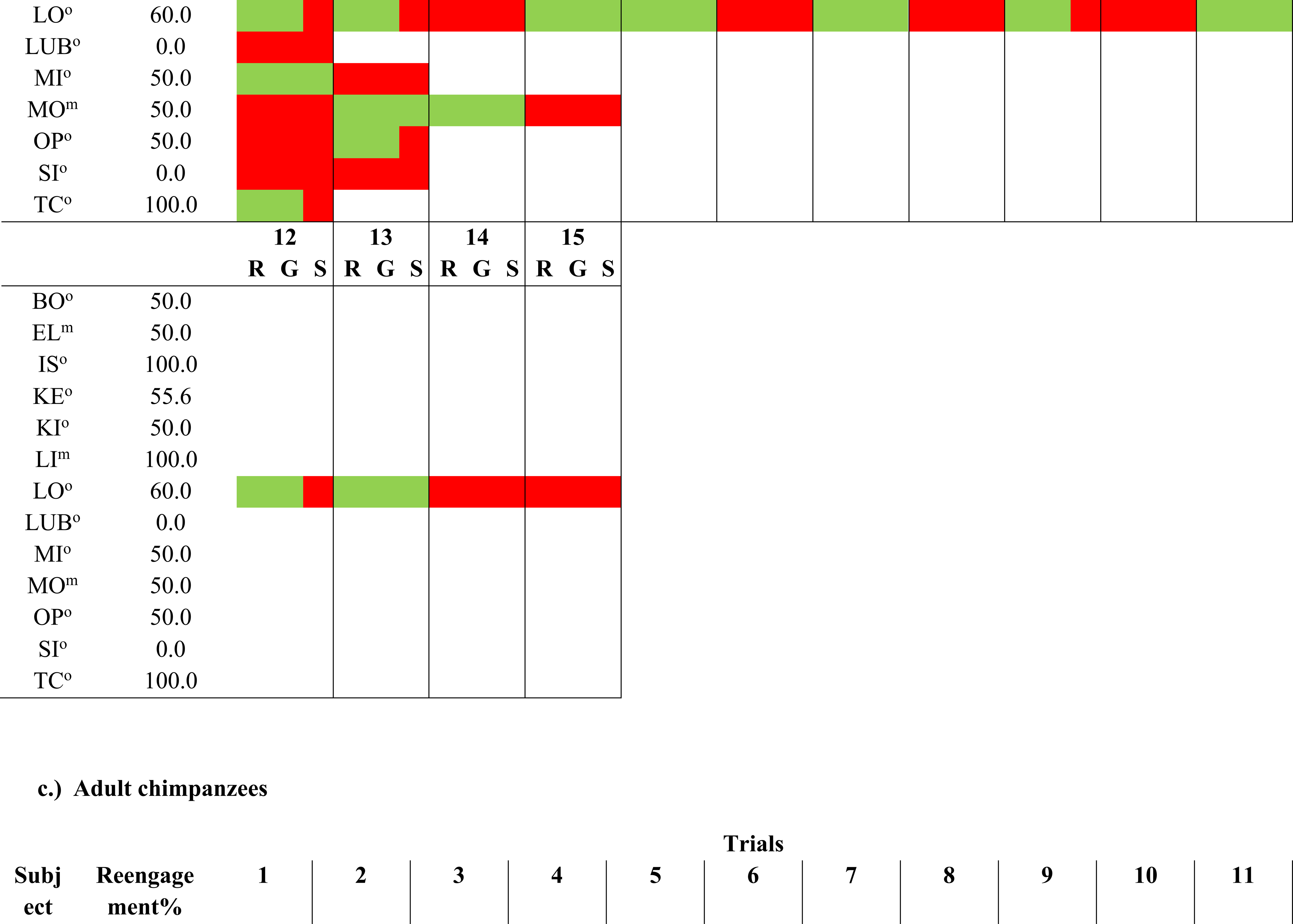
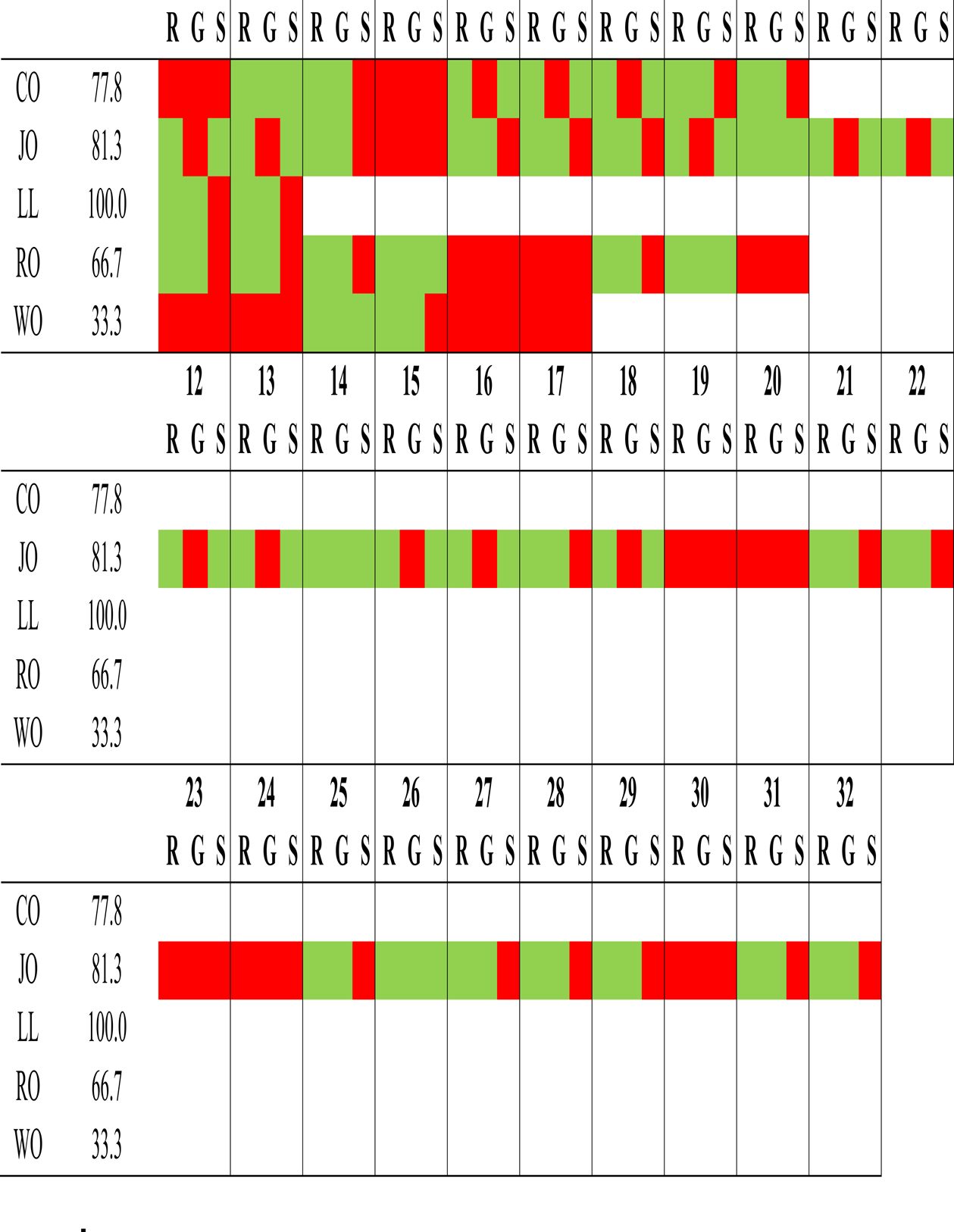
Reengagement, GRB and signal use as per subject and across groups (a: infant bonobos, b: adult bonobos, c: adult chimpanzees). Rearing status for bonobos indicated as ^m^ (mother-reared) and ° (orphan). R = reengagement attempt; G = Use of game related behavior(s); S = Use of signal(s). Red color = Evidence for behavior in question absent; green color = Evidence for behavior in question present. Blank cells indicate absent trials.

When considering reengagement percentages based on all trials (Table 3), we found that in the still-faced condition, infant bonobos attempted to reengage experimenters in six out of seventeen trials (*median* [*IQR*] = 33.3 % [25.0 %; 42.9 %]). In the back-turned condition, they attempted to reengage experimenters in four out of twelve trials (*median* [*IQR*] = 0.0 % [0 %; 50.0 %]). There was no statistical difference in reengagement attempts between conditions (*p* = 0.789).

#### 3.1.2 How do infant bonobos reengage passive partners?

On first trial, none of the infant bonobos used GRBs or signals (0%, Table 3). When considering both first and subsequent trials, one out of five infant bonobos reengaged the experimenter using GRBs (20%) and four out of five infant bonobos did so using signals in at least one trial (80%, Table 3).

Out of all reengaged trials (*N* = 10), we found that only one subject (Kwango, 2.5 y) produced GRBs (i.e., on two out of six reengaged trials). Due to low sample size, we were unable to statistically compare GRB use between conditions for infant bonobos who reengaged. By contrast, infant bonobos used signals in every reengaged trial (*N* = 10), yielding identical signal use across conditions. The signals (*N* = 34) produced by infant bonobos contained 94.1 % gestures, 5.9 % facial expressions and 0 % vocalizations, see S2 Table for raw counts on signal types.

### 3.2 Adult bonobos

#### 3.2.1 Do adult bonobos reengage passive partners?

On first trial, nine out of thirteen adult bonobos reengaged the experimenters (69.2%, Table 3). When considering both first and subsequent trials, eleven out of thirteen adult bonobos reengaged the experimenter in at least in one trial (84.6%, Table 3).

When considering reengagement percentages based on all trials (Table 3), we found that in the still-faced condition, adult bonobos attempted to reengage experimenters in twenty out of thirty trials (*median* [*IQR*] = 66.7 % [56.3 %; 100 %]). In the back-turned condition, they attempted to reengage experimenters in nine out of twenty-two trials (*median* [*IQR*] = 33.3 % [0 %; 100 %]). Despite the variation, there was no statistical difference between conditions (*p* = 0.202).

When looking at rearing status post-hoc, we found that orphans’ and non-orphans’ reengagement percentage was nearly identical, suggesting no direct impact of human scaffolding on reengagement behaviour (orphans *N*=10: median [IQR] = 50.0 % [50 %; 58.9 %]; mother-reared *N*=3: median [IQR] = 50.0 % [50.0 %; 75 %], see Table 3).

#### 3.2.2 How do adult bonobos reengage passive partners?

On first trial, nine out of thirteen adult bonobos used GRB (69.2%) and only two out of thirteen subjects used signals (15.4%), see Table 3. When considering both first and subsequent trials, eleven out of thirteen adult bonobos used GRBs in at least one of their trials (84.6%), while only six out of thirteen subjects produced signals in at least one of their trials (46.2%, Table 3).

Out of all reengaged trials (*N* = 29), the adult bonobos used GRB in twenty-eight of reengaged trials (*median* [*IQR*] = 100 % [100 %; 100 %]). By contrast, they produced signals only in thirteen out of twenty-nine reengaged trials (*median* [*IQR*] = 20 % [0 %; 65.3 %]). The signals (*N* = 26) consisted of 73.1 % gestures, 19.2% facial expressions and 7.7 % vocalizations, see S2 Table for raw counts on signal types. Between conditions, there was no difference in the way bonobos produced GRBs (*p* = 1; still-faced: *median* [*IQR*] = 100 % [100 %; 100 %]; *N* = 20 reengaged trials; back-turned: *median* [*IQR*] = 100 % [100 %; 100 %]; *N* = 9 reengaged trials). Although there were fewer signals produced in the back-turned condition compared to the still-faced condition, the difference in medians likewise did not reach statistical significance (*p* = 1; still-faced: *median* [*IQR*] = 75.0 % [0 %; 100 %]; *N* = 20 reengaged trials; back-turned: *median* [*IQR*] = 0 % [0 %; 18.75 %]; *N* = 9 reengaged trials).

### 3.3 Adult chimpanzees

#### 3.3.1 Do adult chimpanzees reengage passive partners?

On first trial, three out of five adult chimpanzees reengaged the experimenters (60%, Table 3). When considering both first and subsequent trials, all of the five chimpanzees reengaged the experimenters on at least one of their trials (100%, Table 3).

When considering reengagement percentages based on all trials (Table 3), we found that in the still-faced condition, adult chimpanzees attempted to reengage experimenters in twenty-one out of twenty-seven trials (*median* [*IQR*] = 83.3 % [66.7 %; 91.7 %]). In the back-turned condition, they attempted to reengage experimenters in twenty-two out of thirty-one trials (*median* [*IQR*] = 75.0 % [66.7 %; 100 %]). There was no statistical difference between conditions (*p* = 0.584).

#### 3.3.2 How do adult chimpanzees reengage passive partners?

On first trial, two out of five adult chimpanzees used GRB (40%) and only one out of five used signals (20%), see Table 3. When considering both first and subsequent trials, all of the adult chimpanzees used GRBs in at least one of their trials (100%), while four out of five produced signals in at least one of their trials (80%, Table 3).

Out of all reengaged trials (*N* = 43), we found that adult chimpanzees used GRB in thirty reengaged trials (*median* [*IQR*] = 100 % [61.5 %; 100 %]). By contrast, they produced signals only in twenty out of forty-three reengaged trials (*median* [*IQR*] = 50 % [33.3 %; 50.0 %]). The signals (*N* = 26) consisted of 88.5 % gestures, 7.7% facial expressions and 3.8 % vocalizations, see S2 Table for raw signal type counts. Between conditions, there was no difference in the way chimpanzees produced GRBs (*p* = 0.371; still-faced: *median* [*IQR*] = 100 % [54.5 %; 100 %]; *N* = 21 reengaged trials; back-turned: *median* [*IQR*] = 100 % [91.7 %; 100 %]; *N* = 22 reengaged trials). Although there were fewer signals produced in the back-turned condition compared to the still-faced condition, this difference did not reach statistical significance (*p* = 0.181; still-faced: *median* [*IQR*] = 50.0 % [50.0 %; 60.0 %]; *N* = 21 reengaged trials; back-turned: *median* [*IQR*] = 32.5 % [18.8 %; 42.5 %]; *N* = 22 reengaged trials).

### 3.4 Do infant bonobos and adult bonobos differ in terms of reengagement attempts, use of GRBs, or signals?

In the still-faced condition, infant bonobos attempted to reengage experimenters significantly less often compared to adult bonobos (*W =* 45.5, *p* <0.05, *r* = -0.50; Table 3). In the back-turned condition, infant bonobos attempted to reengage the experimenters in fewer trials compared to adult bonobos (Table 3), yet this difference was not statistically significant (*W =* 34.5, *p* = 0.428). In terms of reengagement behaviours, infants used significantly less GRBs compared to adults in reengaged trials (*W =* 44.0, *p* <0.001, *r* = -0.87; Table 3). By contrast, adult bonobos were significantly less likely to use signals compared to infant bonobos in reengaged trials (*W =* 4.0, *p* <0.05, *r* = -0.62; Table 3).

### 3.5 Do adult bonobos and adult chimpanzees differ in terms of reengagement attempts, use of GRBs, or signals?

There were no significant differences in reengagement attempts between adult bonobos and chimpanzees, neither in the still-faced condition (*W =* 25, *p* = 0.817) nor the back-turned condition (*W =* 20.5, *p* = 0.437). Furthermore, there were no significant differences between adult chimpanzees and bonobos in terms of use of GRBs (*W =* 37, *p* = 0.135) or signals (*W =* 25.5, *p* = 0.861) among reengaged trials.

### 3.6 Posthoc analyses to account for variation in the number of trials across subjects

In line with the 30 sec analysis for bonobos, there were no significant differences in the way adult bonobos attempted to reengage experimenters compared to chimpanzees in 15 s interruption periods (still-faced condition: *W =* 18, *p* = 0.303; back-turned condition: *W =* 16.5, *p* = 0.196). There also was no evident correlation between number of trials and reengagement percentages (*rho* = 0.12, *p* = 0.58), indicating no evidence of an increase in reengagement percentages for subjects with greater number of trials.

## 4. Discussion

Reengagement of passive social partners following an interruption of a joint activity has previously been understood as the behavioural indicator of joint commitment [26,30]. Based on contradictory findings regarding reengagement behaviour in apes, our study was designed to expand previous research by assessing whether chimpanzees and bonobos reengage passive social partners in a novel triadic social game (“tug-of-war”). Specifically, our goal was to examine whether the variation in findings across studies regarding apes’ ability to reengage passive partners are affected by subject age, group, and the choice of the game (i.e., the latter being assessed based on comparisons with findings reported by earlier studies). Our data revealed that subjects were motivated to engage in the game with human partners and to reinstate the game when interrupted. Although there were no differences between adult chimpanzee and bonobos, we found differences between infant and adult bonobos, insofar as infant bonobos attempted to reengage partners much less frequently (and never on the first trial) compared to adult bonobos. Interestingly though, when considering all trials, four out of five infant bonobos reengaged the experimenter in at least one of their trials (Table 3), suggesting the ability to reinstate a triadic game in principle. The major difference appeared to lie in the frequency of reengagement across the two age groups, with adult bonobos more readily reengaging compared to infant bonobos. Moreover, when attempting to reengage passive partners, adult bonobos used more GRBs than infants (independent if one considers behaviours on the first trial or all trials), who relied mainly on tactile, gestural communication (without apparent relation to the game) suggesting a better understanding of the joint nature of the game and potential roles of the social partners in adults.

Previous studies have examined reengagement mainly in young individuals [30,34,36], but no consistent within-species comparison has yet been done to assess differences between younger and older age classes. Compared with previous studies, our findings contradicted those of [30], where chimpanzees made no attempts at reengaging a passive experimenter. Instead, our findings add to the growing evidence [24,32,34,36] that apes may possess some of the motivational preconditions to develop an understanding of joint commitment in both dyadic social interactions *and* triadic games. Critically, our current design implied leaving the object accessible to the subjects during interruption periods, which allowed for testing whether the apes would use the game object (garden hose) when attempting to reengage a partner, rather than keeping it to themselves. Indeed, the adult apes in our experiment frequently produced GRBs when reengaging a partner (e.g., handing back the hose, simulating the game action, dropping the hose outside of the cage, touching the experimenter with the hose, see S2 Table), constituting possible evidence of their intentions to resume the game as well as of their understanding of how the game works. The apes’ behaviours may be comparable with the behaviours of human children when engaging in triadic games, who attempt to reengage a passive partner by offering toys or indicating to experimenters how the game was played [e.g., 26].

Although infant chimpanzees have not reengaged passive experimenters in a former study [30], infant bonobos in this study did so, albeit much less compared to adult subjects. Our results thus show that, if conditions are right, even infant apes attempt to reengage experimenters in principle (although less so than adult bonobos and with evidently less complex communicative means). However, given the lack of infant chimpanzees as comparison group, we cannot refute whether the difference between our infant bonobos and the infant chimpanzees tested by [30] could likewise be related to species differences. To test this, future studies should apply this paradigm to reassess reengagement of interrupted triadic joint activities in several groups of infant chimpanzees.

Crucially, although the choice of differential interruption periods across groups (30s for bonobos and 15s for chimpanzees) was intentionally chosen to provide comparable data with previous studies, it represented a limitation for our group comparisons. Yet, an inspection of response latencies across groups (Table 2) showed that adult bonobos’ response latencies were longer than those of chimpanzees. Given this result, we believe that our choice of interruption periods naturally represents the reengagement response latencies of the two groups. The additional posthoc analysis further revealed that when both groups are compared at 15 s, results remain stable (see section 3.6).

When attempting to reengage the experimenter, infants mainly used signals, but very rarely GRBs. Adult bonobos (and chimpanzees), on the contrary, used GRBs on most occasions. Due to the lack of GRBs in infants, it is difficult to ascertain whether they were attempting to reinstate the game. Instead, the infants’ communication might be caused by fear of abandonment (i.e., caretaker turning away). This indicates a more profound understanding of the triadic game in adults as compared to infants; infant bonobos might not yet perceive the interaction as joint in the same degree as older individuals do, a skill that may be scaffolded with social experience and engagement in joint activities. Indeed, infant bonobos have a long development period and stay dependent on their mother until approximately 4-5 years of age [51]. It is possible that reengagement behaviour develops as individuals become more independent in terms of interactions with non-mother partners, gain interactive experience, as well as become more sensitive to social relationships [52,53]. Human children also become skilful at engaging in cooperative activities and reengaging passive partners following their third year of life [23,29,54,55]. Such developmental patterns might explain the negative findings in [30], where juvenile chimpanzees never attempted to reengage experimenters.

One alternative explanation for the increased, especially tactile, gesture use in infant bonobos could be that infants were tested while the experimenter was in their cage, opposite to the adults, where the experimenter was standing outside the cage. If the adult subjects had had the opportunity to touch or grab the experimenter, they might have done so as well. Given the limits of our design, we cannot clarify this here; future research is needed to exclude this explanation. Our findings nonetheless suggest that ontogenetic differences could explain variation in reengagement behaviour and use of GRB across groups. Future research with larger samples and ideally a longitudinal approach would be necessary to solidify the evidence on developmental trajectories of reengagement behaviour in chimpanzees and bonobos.

A potential further factor that might explain differences in reengagement rates between infant and adult bonobos could be the difference in early life and rearing experiences. For instance, one study showed that early manifestations of cooperation varied across two groups of nursery-reared chimpanzees who experienced different caregiving styles in their first year of life [56]. Some of our bonobo subjects are orphans and were raised by human surrogate mothers rather than by their natural mothers (see S2 Table). Such early experiences could have fostered reengagement behaviour, since most of the orphans’ early social experiences have been scaffolded by interactions with humans. However, the difference in rearing experience cannot have had any impact on our results, because none of the apes had any previous experience in playing the tug-of-war games with humans. In support of this view, we found that adult bonobo orphans’ and non-orphans’ reengagement percentage was almost identical, suggesting no direct impact of human scaffolding on reengagement behaviour. Viewed in a different light, one might further argue that a *lack* of knowledge in the tug-of-war game inhibits strong performance, especially in the more wary infants, but this is also unlikely given that subjects’ reengagement attempts did not increase with experience in the game (see section 3).

When considering humans, the impact of the social environment could likewise explain children’s motivation to reengage passive partners. Research on the development of joint commitment in humans is entirely based on WEIRD samples [23,26–29], making it difficult to judge how far empirical manifestations of this capacity are affected by culture and other social dimensions. Given the limited evidence on joint commitment, and small sample sizes, human and ape researchers must be careful in generalizing their findings of some groups to the entire species. More evidence is needed including more samples with varying social and rearing conditions. Moreover, we believe that joint commitment should not only be assessed based on reengagement behaviours, as classically done. Rather, a more inclusive comparative analysis is needed, which examines the way in which participants naturally communicate in spontaneous interactions between peers [14,15,18]. Looking at the process of social action coordination, such as how apes and humans get into and out of interactions [20,21] or how they repair communicative breakdowns [57], could be particularly fruitful to deliver more inclusive and ecologically valid data.

Although our findings, along with previous research [24,32], support the hypothesis that adult bonobos and chimpanzees experience some basic form of joint commitment, it is important to point out that these data do not yet provide conclusive evidence. For instance, apes may simply enjoy playing tug-of-war games and understand that the game requires a cooperating human partner, without any sense of joint commitment. Reengagement behaviour *per se* can thus best be interpreted as a *necessary* but *not sufficient* condition of joint commitment. More certain evidence of joint commitment could be gathered by having apes play in parallel with a human experimenter and compare the reengagement behaviours between social no-commitment and joint commitment conditions [26,38].

Another limitation of our study is that it cannot be determined for certain whether the behaviours observed truly represent reengagement attempts; the current behaviours have been judged as reengagement attempts, but they could likewise represent an initiation attempt to start a new interaction with the experimenter. Apes could equally produce these signals or behaviours in any interaction with humans, potentially even when no game was played. To address this, further baseline controls should be added in the future. For instance, one could add a social control condition to measure the rates by which subjects use these signals and behaviours to communicate with humans in interactions that had *not* been preceded by the tug-of-war game; alternatively, another social baseline control could be one where another type of interaction is reinstated to compare reengagement behaviours following the tug-of-war game and other interaction settings. Though our current research corroborates previous studies showing evidence of reengagement as *behavioural* marker of joint commitment, we acknowledge that more paradigms, controls, and sample groups are needed to determine whether apes have the proclivities to form and understand joint commitments.

Since we tested great apes while engaging with human partners rather than with conspecifics, one could also argue that the apes may not express or develop this quality in their natural environment (i.e., when engaged with conspecifics). Recent results from observational research revealed that apes’ reengagement attempts extend to conspecifics in naturally occurring social grooming and play activities, at least in captivity [24,32]. It remains unclear, however, whether this ability is specific to captive groups or extends to wild apes, which presents an exciting avenue for future research. To address whether reengagement behaviour is specific to *Pan* or *Homo*, further studies might additionally apply this paradigm to other primate species, or animal taxa more distant from humans. One promising recent study has already shown reengagement behaviour in dogs [58], pointing to the possibility of convergent origins of the behavioural correlates of joint commitment.

Although it is often assumed that bonobos are more socially tolerant and pro-social than chimpanzees [41,42], we did not find differences in any of the assessed behaviours between the two groups. In line with another study that compared reengagement rates in natural joint actions of bonobos and chimpanzees [32], our data revealed similar reengagement and signalling rates between the two groups, suggesting that our bonobo subjects were not necessarily more motivated to reconstruct previous commitments with others than our chimpanzee subjects.

Further groups need to be tested to attest proper species differences, something we cannot clarify with this single group comparison. The data nonetheless fits with previous studies on joint commitment, which have shown that chimpanzees, as much as bonobos, appear to exhibit behavioural correlates indicative of their potential engagement in joint commitment [34]. The current findings, along with others [20,24,32], could be seen to point to a continuous evolution of joint commitment, with the early foundation likely having evolved earlier than previously assumed [10], either with (or before) our last common ancestor with *Pan*, or as a convergent adaptation to the demands of social action coordination. Yet, as previously mentioned, further controls need to be placed before firm conclusions can be drawn.

Lastly, in contrast to our prediction, we found no statistical evidence for differences in the reengagement behaviours across conditions in either group. One reason for this may be our small sample size; we were only able to observe trends, notably in adult bonobos, who produced slightly more communicative signals and GRBs when experimenters were facing towards them than when they were facing away. Firm conclusions cannot be drawn from these limited analyses and should be expanded in further studies with more comprehensive sample sizes and conditions of differing intentions of the experimenter. For instance, one might add conditions resembling those used in [59], by having experimenters who are willing to reengage (but unable) or are unwilling to reengage (but able).

## 5. Conclusion

Our findings have shown that chimpanzees and bonobos, even at a young age, have the propensity to reengage a passive partner to a triadic joint game after an interruption. As our data showed, however, infant bonobos communicate less during interruption phases compared to adult bonobos, yielding weaker evidence of reengagement in younger subjects. Primarily, reengagement attempts of young bonobos contained tactile gestures or other signals, while reengagement attempts in adult bonobos (and chimpanzees) often comprised GRBs, indicating a more sophisticated understanding of the joint game in adults. Future studies should attempt to further pinpoint fine-grained differences in behavioural manifestations supporting putative forms of joint commitments in humans and other primates via a bottom-up-approach, investigating all kinds of behaviours and signals, as well as micro bodily movements not necessarily classified as intentional signal. Although reengagement represents one behavioural correlate of joint commitment, we advocate future studies to study additional behavioural correlates of joint commitment, notably signal exchanges to coordinate entries and exit of joint activities [14,15].

## Acknowledgements

We thank La Vallée des Singes and its former director Emmanuel Le Grelle as well as Le Conservatoire pour la Conservation des Primates and ABC Congo, particularly Fanny Minesi, Raphaël Belais and Claudine André, for allowing access to the study sites and subjects. We are particularly thankful for the support of the animal keepers at La Vallée des Singes (Carole Michelet, Franck Alexieff, Raphaël Béguet, Steven Thomas, Thomas Urbain) and at Lola Ya Bonobo (Alain Miteti Muyamba, Patrick Kinkani Kinkani, Ivonne Vela Ntona, C’arrive Nsimba Kiasi, Niclette Bonyoka Mboyo, Micheline Nzozi Katiayi), who acted as experimenters and helped us implement the trials, and to Héritier Izansone, for his invaluable assistance during the experiment.

## Author Contributions

RH, AB, KZ, and EG contributed to conception and design of the study. RH and EG organized the database. RH and KI performed the analysis. RH wrote the first draft of the manuscript. AB, KZ, FR, KI and EG edited and helped revising the manuscript. JG provided resources to ape populations and instructed research at site. All authors contributed to manuscript revision, read, and approved the submitted version.

## Data Availability Statement

The datasets of this study can be found on figshare.com: <u>10.6084/m9.figshare.21175129</u>

## Supporting information captions

**1. Data S1.**

**2. S1 Text.**

**3. S1 Table.** Information about study subjects

**4. S2 Table.** Count of signals and game-related behaviors (GRB) used to reengage the partner across groups

**5. S3 Table.** Definitions of signals (i.e., gestures, vocalizations, facial expressions) and GRBs.

## References

1. Boesch C. Joint cooperative hunting among wild chimpanzees: Taking natural observations seriously. Behav Brain Sci. 2005;28: 692–693. doi:10.1017/S0140525X05230121

2. Holekamp KE, Sakai ST, Lundrigan BL. The spotted hyena (*Crocuta crocuta*) as a model system for study of the evolution of intelligence. J Mammal. 2007;88: 545–554. doi:10.1644/06-MAMM-S-361R1.1

3. Pitman RL, Durban JW. Cooperative hunting behavior, prey selectivity and prey handling by pack ice killer whales (*Orcinus orca*), type B, in Antarctic Peninsula waters. Mar Mammal Sci. 2012;28: 16–36. doi:10.1111/j.1748-7692.2010.00453.x

4. Vail AL, Manica A, Bshary R. Referential gestures in fish collaborative hunting. Nat Commun. 2013;4: 1765. doi:10.1038/ncomms2781

5. Fonio E, Heyman Y, Boczkowski L, Gelblum A, Kosowski A, Korman A, et al. A locally-blazed ant trail achieves efficient collective navigation despite limited information. eLife. 2016;5. doi:10.7554/eLife.20185

6. McCreery HF, Dix ZA, Breed MD, Nagpal R. Collective strategy for obstacle navigation during cooperative transport by ants. J Exp Biol. 2016;219: 3366–3375. doi:10.1242/jeb.143818

7. Clutton-Brock T. Cooperation between non-kin in animal societies. Nature. 2009;462: 51–57. doi:10.1038/nature08366

8. Call J. Contrasting the social cognition of humans and nonhuman apes: The shared intentionality hypothesis. Top Cogn Sci. 2009;1: 368–379. doi:10.1111/j.1756-8765.2009.01025.x

9. Tomasello M, Carpenter M. Shared intentionality. Dev Sci. 2007;10: 121–125. doi:10.1111/j.1467-7687.2007.00573.x

10. Tomasello M. A natural history of human morality. Cambridge, Massachusetts: Harvard University Press; 2016.

11. Clark HH. Social actions, social commitments. In: Enfield NJ, Levinson SC, editors. Roots of human sociality: Culture, cognition and interaction. Oxford, England: Berg; 2006. pp. 126–150.

12. Gilbert M. Joint commitment. In: Jankovic M, Ludwig K, editors. The Routledge handbook of collective intentionality. New York: Routledge; 2017. pp. 130–139.

13. Tomasello M, Moll H. The gap is social: human shared intentionality and culture. In: Kappeler PM, Silk JB, editors. Mind the gap Tracing the origins of human universals. Berlin: Springer; 2010. pp. 331–349.

14. Bangerter A, Genty E, Heesen R, Rossano F, Zuberbühler K. Every product needs a process: unpacking joint commitment as a process across species. Philos Trans R Soc B. 2022;377: 20210095. doi:10.1098/RSTB.2021.0095

15. Heesen R, Genty E, Rossano F, Zuberbühler K, Bangerter A. Social play as joint action: A framework to study the evolution of shared intentionality as an interactional achievement. Learn Behav. 2017;45: 390–405. doi:10.3758/s13420-017-0287-9

16. Leavens DA, Bard KA, Hopkins WD. The mismeasure of ape social cognition. Anim Cogn. 2017. doi:10.1007/s10071-017-1119-1

17. Clark HH. Using language. Cambridge: Cambridge University Press; 1996.

18. Genty E, Heesen R, Rossano F, Zuberbühler K, Guery J, Bangerter A. How apes get into and out of joint actions: precursors of shared intentionality? Interact Stud. 2020;21: 353–386. 10.1075/is.18048.gen

19. Gilbert M. Obligation and joint commitment. Utilitas. 1999;11: 143–163. doi:10.1017/S0953820800002399

20. Heesen R, Bangerter A, Zuberbühler K, Iglesias K, Neumann C, Pajot A, et al. Assessing joint commitment as a process in great apes. iScience. 2021;24: 102872. doi:10.1016/J.ISCI.2021.102872

21. Rossano F, Terwilliger J, Bangerter A, Genty E, Heesen R, Zuberbühler K. How 2- and 4-year-old children coordinate social interactions with peers. Philos Trans R Soc B. 2022;377: 20210100. doi:10.1098/RSTB.2021.0100

22. Chevalley E, Bangerter A. Suspending and reinstating joint activities with dialogue. Discourse Process. 2010;47: 263–291. doi:10.1080/01638530902959935

23. Gräfenhain M, Carpenter M, Tomasello M. Three-year-olds’ understanding of the consequences of joint commitments. PLOS ONE. 2013;8: e73039. doi:10.1371/journal.pone.0073039

24. Heesen R, Bangerter A, Zuberbühler K, Iglesias K, Rossano F, Guéry JP, et al. Bonobos engage in joint commitment. Sci Adv. 2020;6: eabd1306. doi:DOI: 10.1126/sciadv.abd1306

25. Gilbert M. Walking together: A paradigmatic social phenomenon. Midwest Stud Philos. 1990;15: 1–14. doi:10.1111/j.1475-4975.1990.tb00202.x

26. Gräfenhain M, Behne T, Carpenter M, Tomasello M. Young children’s understanding of joint commitments. Dev Psychol. 2009;45: 1430–1443. doi:10.1037/a0016122

27. Kachel U, Svetlova M, Tomasello M. Three-year-olds’ reactions to a partner’s failure to perform her role in a joint commitment. Child Dev. 2017;89: 1–13. doi:10.1111/cdev.12816

28. Kachel U, Svetlova M, Tomasello M. Three- and 5-year-old children’s understanding of how to dissolve a joint commitment. J Exp Child Psychol. 2019;184: 34–47. doi:10.1016/J.JECP.2019.03.008

29. Kachel U, Tomasello M. 3- and 5-year-old children’s adherence to explicit and implicit joint commitments. Dev Psychol. 2019;55: 80–88. doi:10.1037/dev0000632

30. Warneken F, Chen F, Tomasello M. Cooperative activities in young children and chimpanzees. Child Dev. 2006;77: 640–663. doi:10.1111/j.1467-8624.2006.00895.x

31. Tomasello M. Becoming human: A theory of ontogeny. Cambridge, Massachusetts: Harvard University Press; 2019.

32. Heesen R, Zuberbühler K, Bangerter A, Iglesias K, Rossano F, Pajot A, et al. Evidence of joint commitment in great apes’ natural joint actions. R Soc Open Sci. 2021;8: 211121. doi:10.1098/RSOS.211121

33. Hirata S, Fuwa K. Chimpanzees (*Pan troglodytes*) learn to act with other individuals in a cooperative task. Primates. 2007;48: 13–21. doi:10.1007/s10329-006-0022-1

34. MacLean E, Hare B. Spontaneous triadic engagement in bonobos (*Pan paniscus*) and chimpanzees (*Pan troglodytes*). J Comp Psychol. 2013;127: 245–255. doi:10.1037/a0030935

35. Martin CF, Biro D, Matsuzawa T. Chimpanzees spontaneously take turns in a shared serial ordering task. Sci Rep 2017 71. 2017;7: 1–8. doi:10.1038/s41598-017-14393-x

36. Pika S, Zuberbühler K. Social games between bonobos and humans: evidence for shared intentionality? Am J Primatol. 2008;70: 207–210. doi:10.1002/ajp.20469

37. Voinov PV, Call J, Knoblich G, Oshkina M, Allritz M. Chimpanzee Coordination and Potential Communication in a Two-touchscreen Turn-taking Game. Sci Rep 2020 101. 2020;10: 1–13. doi:10.1038/s41598-020-60307-9

38. Tomasello M. The coordination of attention and action in great apes and humans. Philos Trans R Soc B. 2022;377: 20210093. doi:10.1098/RSTB.2021.0093

39. Gruber T. Wild-born Orangutans (Pongo abelii) engage in triadic interactions during play. Int J Primatol. 2014;35: 411–424. doi:10.1007/s10764-013-9745-1

40. Gruber T, Clay Z. A comparison between bonobos and chimpanzees: A review and update. Evol Anthropol Issues News Rev. 2016;25: 239–252. doi:10.1002/evan.21501

41. Hare B, Melis AP, Woods V, Hastings S, Wrangham R. Tolerance allows bonobos to outperform chimpanzees on a cooperative task. Curr Biol. 2007;17: 619–623. doi:10.1016/j.cub.2007.02.040

42. Tan J, Ariely D, Hare B. Bonobos respond prosocially toward members of other groups. Sci Rep. 2017;7: 14733. doi:10.1038/s41598-017-15320-w

43. Jaeggi AV, Stevens JMG, Van Schaik CP. Tolerant food sharing and reciprocity is precluded by despotism among bonobos but not chimpanzees. Am J Phys Anthropol. 2010;143: 41–51.

44. Iki S, Hasegawa T. Face-to-face configuration in Japanese macaques functions as a platform to establish mutual engagement in social play. Anim Cogn 2021 246. 2021;24: 1179–1189. doi:10.1007/S10071-021-01508-1

45. Siposova B, Tomasello M, Carpenter M. Communicative eye contact signals a commitment to cooperate for young children. Cognition. 2018;179: 192–201. doi:10.1016/J.COGNITION.2018.06.010

46. MacLean E, Hare B. Spontaneous triadic engagement in bonobos ({Pan} paniscus) and chimpanzees ({Pan} troglodytes). J Comp Psychol. 2013;127: 245–255. doi:10.1037/a0030935

47. Wittenburg P, Brugman H, Russel A, Klassmann A, Sloetjes H. ELAN: a professional framework for multimodality research. Proceedings of the 5th International Conference on Language Resources and Evaluation (LREC 2006). 2006. pp. 1556–1559.

48. Byrne RW, Cartmill E, Genty E, Graham KE, Hobaiter C, Tanner J. Great ape gestures: intentional communication with a rich set of innate signals. Anim Cogn. 2017;20: 755–769. doi:10.1007/s10071-017-1096-4

49. Landis JR, Koch GG. The measurement of observer agreement for categorical data. Biometrics. 1977;33: 159. doi:10.2307/2529310

50. Field A, Miles J, Field Z. Discovering statistics using R. London: Sage; 2012.

51. Furuichi T, Idani G, Ihobe H, Kuroda S, Kitamura K, Mori A, et al. Population dynamics of wild bonobos Pan paniscus at Wamba. Int J Primatol. 1998;19: 1029–1043. doi:10.1023/A:1020326304074

52. Fröhlich M. Taking turns across channels: Conversation-analytic tools in animal communication. Neurosci Biobehav Rev. 2017;80: 201–209. doi:10.1016/j.neubiorev.2017.05.005

53. Fröhlich M, Wittig RM, Pika S. The ontogeny of intentional communication in chimpanzees in the wild. Dev Sci. 2019;22: e12716. doi:10.1111/desc.12716

54. Eckerman CO, Peterman K. Peers and infant social/communicative development. In: Bremner G, Fogel A, editors. Blackwell handbook of infant development. Oxford, UK: Blackwell Publishing Ltd; 2001. pp. 326–350.

55. Hamann K, Warneken F, Tomasello M. Children’s developing commitments to joint goals. Child Dev. 2012;83: 137–145. doi:10.1111/j.1467-8624.2011.01695.x

56. Bard KA, Bakeman R, Boysen ST, Leavens DA. Emotional engagements predict and enhance social cognition in young chimpanzees. Dev Sci. 2014;17: 682–696. doi:10.1111/desc.12145

57. Heesen R, Fröhlich M, Sievers C, Woensdregt M, Dingemanse M. Coordinating social action: a primer for the cross-species investigation of communicative repair. Philos Trans R Soc B. 2022;377: 20210110. doi:10.1098/RSTB.2021.0110

58. Horschler DJ, Bray EE, Gnanadesikan GE, Byrne M, Levy KM, Kennedy BS, et al. Dogs re-engage human partners when joint social play is interrupted: a behavioural signature of shared intentionality? Anim Behav. 2022;183: 159–168. doi:10.1016/J.ANBEHAV.2021.11.007

59. Call J, Hare B, Carpenter M, Tomasello M. “Unwilling” versus “unable”: chimpanzees’ understanding of human intentional action. Dev Sci. 2004;7: 488–498. doi:10.1111/j.1467-7687.2004.00368.x

