## Supplementary information for "Chimpanzees and bonobos reinstate an interrupted triadic game"

### 1 S1 Data

The datasets of this study can be found on figshare.com: 10.6084/m9.figshare.21175129

### 2 S1 Text

We additionally attempted two conditions, which were randomly administered alongside other conditions. These conditions had to be discarded post-hoc from the main analyses due to methodological limitations.

In the third attempted condition ("clumsy request"), conducted in all groups, the experimenter was supposed to simulate being clumsy, by reaching to the hose with the hand but not being "able" to properly grab it. The problem was that the hose could either stick out through the cage mesh, reachable for the experimenter, or it could fall within the cage, inaccessible for the experimenter. The clumsy request relied on different intentions in these two cases; in the first, the experimenter seems willing but unskilful (but could potentially grab the hose), and in the second, the experimenter seems willing but incapable (could never grab the hose even if willing to do so).

In a fourth attempted condition ("third-party interruption"), only applied to adult and infant bonobos, the experimenter was interrupted by another person, who approached and started to talk to the experimenter. This condition was only implemented in bonobos and contained errors due to difficulty in implementation and a very small sample size; it shall be repeated by standardizing experimenter's behaviours, since the experimenters often made mistakes, leading to inconsistencies in the test. For instance, they were either a) turning too slowly toward the third person who interrupts, b) holding the hose for too long while already starting to talk to the third person, c) not engaging fully in the conversation with third person, or d) turning around to check, due to other conflicts in the group of bonobos, or other disturbing bonobos.

**S1 Table.** Information about study subjects. Ages for infant bonobos were estimated, as precise birthdates of orphans were unknown. Mo.-reared = Mother-reared.

| Subject (abbreviated ID) | Species | Site | Age (years) | Age category (this study) | Age class (Kano, 1992) | Sex | Rearing (for sanctuary bonobos) |
|---|---|---|---|---|---|---|---|
| BA | Bonobo | Lola Ya Bonobo | 3.0 | Infant | Infant | Male | Orphan |
| BI | Bonobo | Lola Ya Bonobo | 4.0 | Infant | Infant | Male | Orphan |
| KW | Bonobo | Lola Ya Bonobo | 2.5 | Infant | Infant | Male | Orphan |
| LA | Bonobo | Lola Ya Bonobo | 4.5 | Infant | Infant | Female | Orphan |
| LU | Bonobo | Lola Ya Bonobo | 2.5 | Infant | Infant | Female | Orphan |
| BO | Bonobo | Lola Ya Bonobo | 9.0 | Adult | Subadult | Female | Orphan |
| EL | Bonobo | Lola Ya Bonobo | 6.0 | Adult | Juvenile | Female | Mo.-reared |
| IS | Bonobo | Lola Ya Bonobo | 21.0 | Adult | Adult | Female | Orphan |
| KE | Bonobo | Lola Ya Bonobo | 25.0 | Adult | Adult | Male | Orphan |
| KI | Bonobo | Lola Ya Bonobo | 8.0 | Adult | Subadult | Female | Orphan |
| LI | Bonobo | Lola Ya Bonobo | 8.0 | Adult | Subadult | Female | Mo.-reared |
| LO | Bonobo | Lola Ya Bonobo | 11.0 | Adult | Subadult | Male | Orphan |
| LUB | Bonobo | Lola Ya Bonobo | 6.0 | Adult | Juvenile | Female | Orphan |
| MI | Bonobo | Lola Ya Bonobo | 8.0 | Adult | Subadult | Female | Orphan |
| MO | Bonobo | Lola Ya Bonobo | 11.0 | Adult | Subadult | Male | Mo.-reared |
| OP | Bonobo | Lola Ya Bonobo | 23.0 | Adult | Adult | Female | Orphan |
| SI | Bonobo | Lola Ya Bonobo | 10.0 | Adult | Subadult | Male | Orphan |
| TC | Bonobo | Lola Ya Bonobo | 30.0 | Adult | Adult | Female | Orphan |
| CO | Chimpanzees | La Vallée des Singes | 22.0 | Adult | Adult | Male | - |
| JO | Chimpanzees | La Vallée des Singes | 23.0 | Adult | Adult | Male | - |
| LL | Chimpanzees | La Vallée des Singes | 9.0 | Adult | Subadult | Female | - |
| RO | Chimpanzees | La Vallée des Singes | 21.0 | Adult | Adult | Male | - |
| WO | Chimpanzees | La Vallée des Singes | 21.0 | Adult | Adult | Male | - |

**S2 Table.** Count of signals and game-related behaviors (GRB) used to reengage the partner across groups.

| Reengagement type | Signal/GRB type | Infant bonobos | Adult bonobos | Adult chimpanzees |
|---|---|---|---|---|
| Gesture | Bang cage | 0 | 0 | 1 |
| Gesture | Bipedal swagger | 0 | 0 | 2 |
| Gesture | Dangle | 2 | 0 | 0 |
| Gesture | Grab experimenter | 5 | 0 | 0 |
| Gesture | Hit object | 1 | 0 | 0 |
| Gesture | Hit object with object | 1 | 4 | 1 |
| Gesture | Jump | 1 | 0 | 0 |
| Gesture | Knock mesh | 0 | 0 | 2 |
| Gesture | Move hose | 3 | 0 | 0 |
| Gesture | Present body | 5 | 1 | 1 |
| Gesture | Present genitals | 1 | 2 | 0 |
| Gesture | Reach | 4 | 0 | 4 |
| Gesture | Shake experimenter | 2 | 0 | 0 |
| Gesture | Shake hand | 0 | 2 | 0 |
| Gesture | Shake head | 1 | 1 | 1 |
| Gesture | Shake hose | 3 | 8 | 5 |
| Gesture | Stomp | 0 | 0 | 3 |
| Gesture | Touch experimenter | 3 | 0 | 1 |
| Gesture | Touch hose | 0 | 1 | 2 |
| GRB | Hand back hose | 1 | 52 | 13 |
| GRB | Drop hose out of cage | 0 | 6 | 7 |
| GRB | Prompting | 0 | 5 | 6 |
| GRB | Simulate game action | 0 | 3 | 10 |
| GRB | Touch experimenter with hose | 1 | 2 | 9 |
| Facial expression | Play face | 2 | 3 | 0 |
| Facial expression | Pout face | 0 | 2 | 1 |
| Facial expression | Tightened lips | 0 | 0 | 1 |
| Vocalization | Hoo | 0 | 0 | 1 |
| Vocalization | Laughter | 0 | 1 | 0 |
| Vocalization | Peep | 0 | 1 | 0 |

**S3 Table.** Definitions of signals (i.e., gestures, vocalizations, facial expressions) and GRBs (after Byrne et al., 2017; Clay et al., 2015; Crockford et al., 2018; de Waal, 1988; Genty et al., 2014, 2015).

| Type | Description |
|---|---|
| **Gestures** | |
| Bang cage | Hand(s) (or feet) brought into hard audible contact with the cage mesh (can be repetitive) |
| Bipedal swagger | Side to side of forward and back movement while standing/walking bipedal (rarely also quadrupedal) |
| Dangle | Subject hangs from arm(s) from the top of the cage/mesh, with feet shaking loosely |
| Grab experimenter | Hand(s) firmly closed over part of experimenter's body |
| Hit object | Hand(s) brought into short hard contact with the surface of an object |
| Hit object with object | Object brought into short hard contact with another object |
| Jump | While in bipedal position both feet leave ground simultaneously with horizontal displacement |
| Knock mesh/object | Back of hand/knuckles brought into short hard audible contact with cage mesh/object (can be repetitive) |
| Move hose | Hose is displaced in one direction, contact with hose is maintained throughout |
| Present body | Body part moved to deliberately expose an area to experimenter's attention |
| Present genitals | Genitals moved to deliberately expose them to experimenter's attention |
| Reach | Arm is extended to the experimenter with hand in open, palm exposed position |
| Shake experimenter | Hand(s) is firmly closed over part of experimenter's body with repeated back and forth motion of hand |
| Shake head | Head moved back and forth in small, repeated side motion |
| Shake hose | Hand(s) firmly closed over hose with repeated up and down or side movements |
| Shake hand | Small repeated back and forth motion of hand(s) from wrist |
| Stomp | Sole of the foot/feet brought into short hard audible contact with surface |
| Touch experimenter | Palm of hand and/or body is brought to light contact with part of experimenter's body, for under 2 s. Also counted if subject attempts to touch experimenter through the mesh but fingers are too short to reach experimenter's body |
| Touch hose | Palm of hand and/or body is brought to light contact with hose, for under 2 s |
| **GRBs** | |
| Drop hose out of cage | Hose pushed through the mesh outside of cage and out of reach (from subject) |
| Hand back hose | One end of hose pushed through the cage mesh outside of cage, while hand is firmly closed over other end of hose |
| Prompt | Body/hand is moved in rapid, short, and tense movements, while hand is firmly closed over hose and while looking at experimenter |
| Simulate game action | Hose pushed back and forth through the mesh (mimicking the game's pull and relax movements) |

| | |
|---|---|
| Touch with hose | Hose pushed through the cage mesh outside of cage and brought to light contact over part of experimenter's body |
| **Vocalizations** | |
| Laughter | Voiceless breathing sounding like low-pitched grunting |
| Hoo | Quiet, relatively inconspicuous vocalization sounding like «*hoo*», mostly emitted as a single call |
| Peep | High-frequency, often closed-mouth vocalization; short in duration and flat, simple acoustic structure |
| **Facial expressions** | |
| Pout face | Lips pursed forward and curled outward in front (circular opening); lips pressed together at mouth corners |
| Tightened lips | Lips horizontally tensed; drawn inward or slightly baring teeth |
| Play face | Mouth opened with lips either in relaxed position covering the upper teeth completely and the lower teeth partially, or retracted, showing both upper and lower frontal teeth |
